# Magicplex^TM^ Sepsis real-time test for the rapid diagnosis of bloodstream infections in adults

**DOI:** 10.1101/374736

**Authors:** Yuliya Zboromyrska, Catia Cillóniz, Nazaret Cobos-Trigueros, Manel Almela, Juan Carlos Hurtado, Andrea Vergara, Caterina Mata, Alex Soriano, Josep Mensa, Francesc Marco, Jordi Vila

## Abstract

Sepsis is a serious health condition worldwide, affecting more than 30 million people globally each year. In 2017, the World Health Organization (WHO) adopted a resolution on sepsis: “improving the prevention, diagnosis and clinical management of sepsis”, with the aim of improving early diagnosis, management and prevention to save lives. Blood culture (BC) is generally used to diagnose sepsis because of the low quantity of microbes occurring in the blood during such infections. However, approximately 50% of bloodstream infections (BSI) give negative BC, this figure being higher for sepsis, which delays the start of appropriate antimicrobial therapy. This prospective study evaluated a multiplex real-time polymerase chain reaction, the Magicplex^TM^ Sepsis test (MP), for the detection of pathogens from whole blood, comparing it to routine BC. We analysed 809 blood samples from 636 adult patients, with 132/809 (16.3%) of the samples positive for one or more relevant microorganism according to BC and/or MP. The sensitivity and specificity of MP were 29% and 95%, respectively, while the level of agreement between BC and MP was 87%. The rate of contaminated samples was higher for BC (10%) than MP (4.8%) (*P* < 0.001). Patients with only MP-positive samples were more likely to be on antimicrobial treatment (47%) than those with only BC-positive samples (18%) (*P* = 0.002). In summary, the MP test reduces the time taken to identify the microbial pathogen, improving the diagnosis of BSI in patients on antibiotic treatment.

## Introduction

In 2017, the World Health Organization (WHO) adopted a resolution on sepsis: “improving the prevention, diagnosis and clinical management of sepsis”, with the aim of improving early diagnosis, management and prevention to save lives. Blood culture (BC) is generally used to diagnose sepsis because of the low quantity of microbes occurring in the blood during such infections. However, approximately 50% of bloodstream infections (BSI) give negative BC, this figure being higher for sepsis, which delays the start of appropriate antimicrobial therapy and consequently results in worse outcomes and higher mortality rates (1,2). Prompt microbiological diagnosis enables a more appropriate antimicrobial treatment than the empirical combination of broad-spectrum antibiotics that have negative effects such as an increased prevalence of resistant pathogens (3).

The microbial diagnosis of sepsis by BC has two main advantages. First, it allows the growth of very small numbers of the microorganism, which is important since the concentration of bacteria in the blood of septic adult patients is usually low (< 10 CFU/mL) (4). Second, this technique allows the isolation of pathogens and, hence, antimicrobial susceptibility testing can be performed. However, BC also has limitations that do not make it an ideal gold standard test, including the long time required for growth detection, the frequent false negative results in patients receiving antimicrobial therapy, and the inability to detect fastidious microorganisms (5). Molecular tests offer important advantages over BC that could improve the diagnosis of BSI, such as the lower amount of time taken to obtain results by working directly from blood. In addition, the low detection limits of molecular assays might make them more sensitive than BC, enabling the detection of fastidious, non-viable or non-culturable microorganisms, even from patients on antibiotic treatment (5–7). Another advantage of molecular assays is the ability to detect some specific resistance markers, which can provide important information for better treatment. Moreover, the rapid identification of the microorganism can be used to infer antimicrobial susceptibility according to local epidemiology.

The Magicplex^TM^ Sepsis test (MP) (Seegene, Seoul, South Korea) is a multiplex real-time polymerase chain reaction (PCR) that detects more than 90 microorganisms at the genus level (73 Gram-positive bacteria, 12 Gram-negative bacteria and 6 fungi), 27 at the species level and 3 drug-resistant genes (*mecA*, *vanA* and *vanB)* within 6 hours. In this study, we evaluated the ability of the MP test to rapidly detect pathogens causing BSI in adult patients from whole blood compared to conventional BC.

## Methods

### Setting, data and sample collection

Paired BC and 1 ml samples of whole blood in an EDTA tube were obtained from adult patients (≥ 18 years old) from the Hospital Clinic of Barcelona, a 700-bed university hospital in Barcelona, Spain, from May to September 2011.

Samples were obtained from patients with suspected BSI and who met the criteria for BC collection. Samples were processed in parallel by MP and BC. Blood samples for MP and BC were obtained simultaneously from the same catheter or venipuncture. For each case, additional clinical data about ongoing antibiotic treatment, the suspected source of infection, as well as the results of other microbiological tests were collected. The study was approved by the Ethics Committee of the Hospital Clinic of Barcelona (study no. 2011/6613).

### Routine microbiological techniques

BC was incubated in Bactec 9240^®^ (Becton Dickinson, MD, USA) for a maximum of 5 days. The following vials were used: the resin-containing BACTEC Plus Aerobic/F and BACTEC Plus Anaerobic/F, and the non-resin-containing BACTEC Standard/10 Aerobic/F and BACTEC Lytic/10 Anaerobic/F. For positive samples, Gram staining and culturing on solid media were performed. Microorganisms were identified using matrix-assisted laser desorption/ionization time-of-flight mass spectrometry (MALDI-TOF MS) (Bruker Daltonics, Bremen, Germany). Routine susceptibility testing included the Phoenix^TM^ system (Becton Dickinson, MD, USA), Etest (AB bioMérieux), microdilution (Sensititre, Trek Diagnostic Systems, Inc., Westlake, Ohio, USA) and the disc diffusion method, depending on the pathogen isolated. Determination of the isolates as contaminants or pathogens was achieved by combining the clinical setting, pathogenicity of the isolated microorganisms and the number of positive BC vials in the case of potential skin contaminants, such as coagulase-negative staphylococci (CoNS). CoNS were considered pathogens if the same species was detected in both sets of BC showing the same antimicrobial susceptibility pattern. Samples with positive BC for pathogens not included in the MP panel were excluded from the analysis.

### The Magicplex^TM^ Sepsis test (MP)

Blood samples were initially pre-treated with the Seegene Blood Pathogen Kit^TM^, according to the manufacturer’s instructions. Bacterial DNA was automatically extracted and purified with the SeePrep12^TM^ extractor (Seegene). Amplification, screening and identification were performed following the manufacturer’s instructions. Each sample was first processed by two conventional PCRs performed in two separate tubes: one for the detection of Gram-positive bacteria and resistance genes, and the other for Gram-negative bacteria (GNB) and fungi (Fig 1). The first real-time PCR (rt-PCR) for screening was performed in three separate tubes, allowing the identification of pathogens at the genus or group level: *Streptococcus* spp., *Staphylococcus* spp., *Enterococcus* spp., GNB group A or B, fungi and 3 resistance genes (*vanA*, *vanB* and *mecA*). For positive results, the second rt-PCR was carried out, allowing the identification of 27 pathogens at the species level, including 3 *Streptococcus* spp. (*Streptococcus pneumoniae*, *Streptococcus agalactiae* and *Streptococcus pyogenes*), 3 *Enterococcus* spp. (*Enterococcus faecalis*, *Enterococcus faecium* and *Enterococcus gallinarum*), 3 *Staphylococcus* spp. (*Staphylococcus epidermidis*, *Staphylococcus haemolyticus* and *Staphylococcus aureus*), 6 GNB from group A (*Pseudomonas aeruginosa*, *Acinetobacter baumannii*, *Stenotrophomonas maltophilia*, *Serratia marcescens*, *Bacteroides fragilis* and *Salmonella typhi*), 6 GNB from group B (*Klebsiella pneumoniae*, *Klebsiella oxytoca*, *Proteus mirabilis*, *Escherichia coli*, *Enterobacter cloacae* and *Enterobacter aerogenes*), and 6 fungi (*Candida albicans*, *Candida parapsilosis*, *Candida glabrata*, *Candida tropicalis*, *Candida krusei* and *Aspergillus fumigatus*).

**Figure 1.**
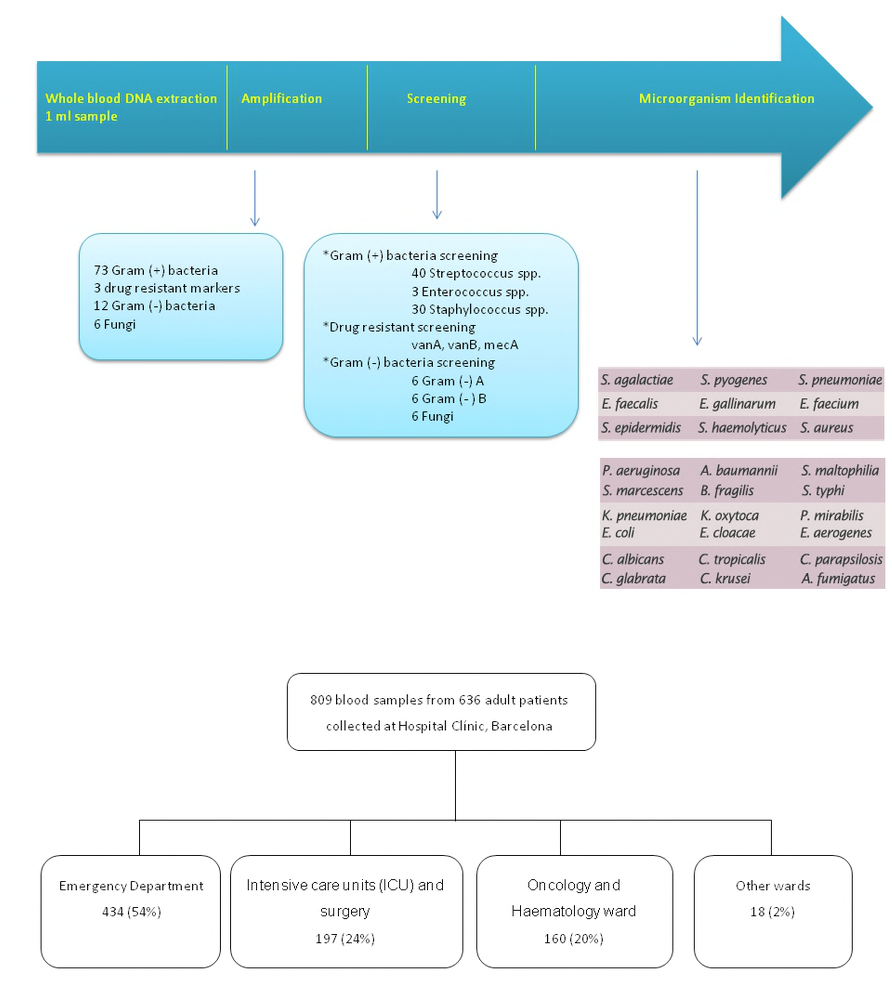
Process of Magicplex ^TM^ Sepsis.

The Seegene Viewer software was used to interpret the results, which were available within 6 hours. All the samples were stored at 2-8°C and processed within 24 h after collection.

MP samples positive for CoNS were considered to be true positives if the same microorganism was detected in two sets of BC. Samples that were positive according to MP in the first rt-PCR screening and identified only at the group level (e.g., GNB group A), but negative in the second rt-PCR were considered negative according to MP. Microorganisms identified by MP only at the genus level, such as *Staphylococcus* spp. and *Streptococcus* spp., were considered to be contaminants if the BC was negative or positive for staphylococci or streptococci that could be identified by MP at the species level.

### Statistical analysis

Statistical analyses were performed using the SPSS software (SPSS, Chicago, IL, USA). Differences were considered significant when *P* < 0.05.

## Results

A total of 809 samples from 636 adult patients were collected and analysed. Among the samples, 434 (54%) were from the emergency wards, 197 (24%) from intensive care units (ICUs) and surgery, 160 (20%) from oncology and haematology wards, and 18 (2%) from other medical wards (Fig 2).

A total of 140 pathogens were detected among 132 positive samples (Table 1). *E. coli* was the most frequently detected pathogen (47/140, 33.6%), followed by CoNS (16/140, 11.4%), *Candida* spp. (14/140, 10%) and *S. aureus* (13/140, 9.3%).

**Table 1.** Comparison of microbiological results between BC and MP.

| Pathogens Detected | Positive Only BC | Positive Only MP | Positive Both Methods | Total , n (%) |
| --- | --- | --- | --- | --- |
| *Escherichia coli* | 27 | 15 | 5 | 47 (34) |
| CoNS | 7 | 0 | 9 | 16 (11) |
| *Candida* spp. | 9 | 2 | 3 | 14 (10) |
| *Staphylococcus aureus* | 2 | 7 | 4 | 13 (9) |
| *Enterococcus faecalis* | 5 | 3 | 3 | 11 (8) |
| *Pseudomonas aeruginosa* | 6 | 2 | 1 | 9 (6) |
| *Klebsiella pneumoniae* | 6 | 2 | 1 | 9 (6) |
| *Streptococcus* spp. | 3<sup>a</sup> | 2<sup>b</sup> | 1<sup>c</sup> | 6 (4) |
| *Enterobacter cloacae* | 3 | 1 | 0 | 4 (3) |
| *Enterococcus faecium* | 2 | 0 | 1 | 3 (2) |
| *Acinetobacter* spp. | 1 | 2 | 0 | 3 (2) |
| *Serratia marcescens* | 1 | 1 | 0 | 2 (2) |
| *Enterococcus gallinarum* | 0 | 1 | 0 | 1 (1) |
| *Klebsiella oxytoca* | 0 | 1 | 0 | 1 (1) |
| *Stenotrophomonas maltophilia* | 0 | 1 | 0 | 1 (1) |
| Total, n(%) | 72 (51) | 40 (29) | 28 (20) | 140 (100) |

BC, blood culture; MP, MagicPlex assay; CoNs, coagulase-negative staphylococci a Streptococci of viridians group (*S. anginosus* and *S. gordonii*) and one *S. agalactiae* b One *S. agalactiae* and one *S. pneumoniae* c *S. constellatus*

BC identified 3 mixed infections: *E. cloacae* plus *K. pneumonia*e, *C. glabrata* plus *S. haemolyticus* and *P. aeruginosa* plus *S. epidermidis*. MP detected 4 mixed infections: *S. agalactiae* plus *S. aureus*, *E. coli* plus *S. epidermidis*, *P. aeruginosa* plus *S. marcescens* and *E. coli* plus *E. gallinarum*. One mixed infection (*E. faecium* plus *S. haemolyticus*) was identified by both methods.

Three *mecA* genes were detected among the 9 pathogenic CoNS identified by the two methods. Routine susceptibility testing detected 7 methicillin-resistant CoNS and 2 methicillin-susceptible strains. Only one *vanB* resistance marker was detected in a sample positive for *E. faecalis* only according to MP and not BC. Therefore, this result remained unconfirmed. No additional BC or other microbiological samples positive for *E. faecalis* were obtained from this patient.

As shown in Table 2, samples with microbial contaminants were more frequently observed with BC than MP: 81/809 (10%) versus 39/809 (4.8%) (*P* < 0.001), respectively.

**Table 2.** Concordance of the results obtained by BC and MP.

|  | Number of Samples |  |  |  |
| --- | --- | --- | --- | --- |
|  | BC positive | BC negative | BC-contaminated | Total |
| **MP-positive** | 28 | 31 | 5 | 64 |
| **MP-negative** | 62 | 574 | 70 | 706 |
| **MP-contaminated** | 6 | 27 | 6 | 39 |
| **Total** | 96 | 632 | 81 | 809 |

BC, blood culture; MP, MagicPlex assay

Blood samples in which only microbial contaminants were detected were not considered for further analysis. As a result, 132/809 (16.3%) of the samples were considered positive for one or more relevant microorganism by BC and/or MP. The level of agreement between BC and MP was 87.1% (705/809). Regarding the 28 samples positive according to both methods, the same pathogen was identified in 27 of the cases, with the remaining one sample showing inconsistent results: *E. coli* detected by MP and *E. faecium* by BC. The rate of positive results was higher for BC (96/809, 11.9%) than MP (64/809, 7.9%). As BC is considered the gold standard, the sensitivity, specificity, positive predictive value and negative predictive value of MP were 29.2% (95% CI, 20.6-39.5), 95% (95% CI, 93-96.4), 43.8% (95% CI, 31.6-56.7) and 90.9% (95% CI, 88.5-92.8), respectively.

Additional clinical information, including underlying disease and the antibiotic or antifungal therapy administered on the day of sampling, was recorded for the patients with 36 MP-positive and BC-negative or contaminated results (Table 3). When comparing the effects of the antimicrobial treatment at the time of sample collection among the three groups of patients with positive samples (only BC-positive (12/68, 17.6%), only MP-positive (17/36, 47.2%), and BC and MP-positive (8/28, 28.6%)), a statistically significant difference was found between the group with only MP-positive samples and that with only BC-positive samples (*P* = 0.002). Interestingly, 3 of the 17 patients with only an MP-positive sample had a BC positive for the same microorganism, while in 5 patients, other microbiological samples were positive for the pathogen detected by MP.

**Table 3.** Additional information from 36 patients with MP-positive and BC-negative results.

| Setting | MP result | Clinical condition | Other microbiological tests positive for the same pathogen | Ongoing antibiotic/antifungal treatment<sup>a</sup> |
| --- | --- | --- | --- | --- |
| ICU and Surgery (n=16) | *Klebsiella oxytoca* | Self-limiting fever after removal of a Kehrs tube in a liver transplant patient | - | - |
|  | *Escherichia coli* | Subarachnoid hemorrhage<br>Control BC 3 days after catheter-related<br>*Pseudomonas aeruginosa* bacteremia | - | Ciprofloxacin |
|  | *Escherichia coli* | Necrotizing fasciitis | BC and wound swab culture positive for *E. coli* 3 days before sampling | Meropenem |
|  | *Staphylococcus aureus* | Necrotizing fasciitis | BC positive for *S. aureus* 2 days before sampling | Cloxacillin |
|  | *Streptococcus pneumoniae* | Acute myeloid leukemia with neutropenic fever without a clinical focus | - | Piperacillin-tazobactam, Vancomycin |
|  | *Candida parapsilosis* | *K. pneumoniae* bacteremia secondary to mesenteric vein thrombosis in a cirrhotic patient<br>Nosocomial pneumonia due to *S.aureus* and *Acinetobacter baumannii* | 3 bronchial washing samples positive for *Candida* spp. | Fluconazole |
|  | *Escherichia coli* | Septic shock due to *S. aureus* cellulitis in a cirrhotic patient. | Wound smear positive for *S. aureus* | Ceftazidime and Tigecycline |
|  |  | Ascites decompensation |  |  |
|  | *Escherichia coli* | Acute necrotizing pancreatitis with peripancreatic abscess | - | Meropenem |
|  | *Staphylococcus aureus* | Superinfection of Pulmonary contusion in a polytrauma patient | Tracheal aspirate positive for *S. aureus* two days before sampling | Levofloxacin |
|  | *Escherichia coli* | Colonic perforation post-nephrectomy | - | Piperacillin-tazobactam |
|  | *Enterobacter cloacae* | Intestinal obstruction and abdominal sepsis secondary to inguinal hernia | - | Levofloxacin |
|  | *Acinetobacter baumannii* | Postoperative fever (liver resection 48 h before sampling) | - | - |
|  | *Acinetobacter baumannii* | Thoracic empyema in a cirrhotic patient | - | - |
|  | *Staphylococcus aureus* | Hepatocellular carcinoma. Metabolic decompensation of diabetes mellitus | - | - |
|  | *Escherichia coli* | Urinary tract infection (UTI) | - | Ciprofloxacin and Ceftriaxone |
|  | *Candida parapsilosis* | Postoperative intraabdominal abscess | - | - |
| **Emergency Department (n=15)** | *Escherichia coli* | UTI | - | - |
|  | *Staphylococcus aureus* | Stroke. Bronchoaspiration in a patient with prior colonization with *S. aureus*. | - | - |
|  | *Streptococcus agalactiae*, | Cellulitis | - | - |
|  | *Staphylococcus aureus* |  |  |  |
|  | *Escherichia coli* | HIV patient with prosthetic aortic valve with progressive prurigo nodularis | - | Amoxicillin/clavulanic acid |
|  | *Escherichia coli* | Fever during hemodialysis | Urine culture positive for *E. coli* | Amoxicillin/clavulanic acid |
|  | *Escherichia coli* | Acute exacerbation of chronic obstructive pulmonary disease (COPD) | Sputum sample positive for *E. coli* | - |
|  | *Enterococcus faecalis* | Traveller's diarrhea | - | - |
|  | *Pseudomonas aeruginosa* | Acute gastroenteritis and community-acquired UTI | - | - |
|  | *Staphylococcus aureus* | Pneumonia | - | - |
|  | *Staphylococcus aureus* | UTI in a patient with permanent bladder catheter | - | Amoxicillin/clavulanic acid |
|  | *Pseudomonas aeruginosa* | Acute exacerbation of Crohn's disease. | - | - |
|  | *Klebsiella pneumoniae* | *Pneumocystis carinii* pneumonia in an AIDS patient | - | - |
|  | *Stenotrophomonas maltophilia* | Community-acquired UTI | - | - |
|  | *Escherichia coli* | Acute exacerbation of COPD | - | - |
|  | *Klebsiella pneumoniae* | *Pneumocystis carinii* pneumonia in an AIDS patient. Bacteremia caused by *E. faecalis* | - | - |
| **Oncology and** |  |  |  |  |

| Haematology (n=4) |  |  |  |  |
| --- | --- | --- | --- | --- |
|  | *Enterococcus faecalis*<br>(vanB positive) | Non-Hodgkin T-cell lymphoma.<br>Neutropenic fever | - | Levofloxacin |
|  | *Escherichia coli* | Acute myeloid leukemia.<br>Neutropenic fever | - | - |
|  | *Escherichia coli* | Stem cell transplanted patient<br>due to acute myeloid leukemia.<br>Neutropenic fever | - | - |
|  | *Escherichia coli* | Fever in a patient with acute<br>lymphoblastic leukemia | - | Meropenem |
|  | *Enterococcus faecalis* | Catheter-related blood stream<br>infection in a patient<br>transplanted due to acute<br>myeloid leukemia | BC positive for *E. faecalis* 6 days<br>before sampling. | Daptomycin |

ICU: intensive care unit; BC, blood culture; MP, MagicPlex assay
<sup>a</sup> Only antimicrobial/antifungal therapy potentially effective against pathogen detected and initiated before sampling

## Discussion

Several molecular assays have been recently tested for the direct molecular identification of pathogens in blood samples (8–13). However, few studies have focused on commercially available PCR-based tests other than the SeptiFast test for the detection of BSI (14,15). In the present study, we evaluated the MagicplexTM Sepsis test (MP), comparing it with conventional BC. We observed that MP showed a sensitivity and specificity of 29.2% and 95%, respectively. Only a few studies have evaluated the sensitivity and specificity of MP (10, 16–19). Carrara et al. (16), assessing 267 patients from ICU, emergency and haematology wards, reported an overall sensitivity of 65% for MP compared with 41% for BC. However, the authors calculated the sensitivities and specificities using a reference standard in which clinical data and other cultures were added to the BC results to see if a positive MP result represented a true BSI or not. Loonen et al. (10), investigating 125 patients from the emergency department, reported a sensitivity of 37% and a positive predictive value (PPV) of 30% for MP, the low PPV resulting from many of the samples being positive for CoNS according to MP. Ljungstrom et al. (17), analysing samples from 375 patients with suspected sepsis collected from emergency wards, excluded suspected contaminants such as CoNS. They reported a sensitivity of 64% and specificity of 96% for MP. However, if they had included all the MP and BC results, the sensitivity of MP would have been 38%, PPV 17% and specificity 65%. Taking into account all these results, we conclude that MP can be useful for the rapid detection of pathogens causing sepsis.

The low sensitivity of MP is probably due to the low bacterial concentration in whole blood and the low sample volume of 1 ml used. There are several strategies that can be applied to improve bacterial recovery, such as removing or significantly reducing the amount of human DNA present in the whole blood sample. This step is included in the MP and SepsiTest assays. Another possibility is to introduce an additional incubation step prior to extraction (20). Although this approach can increase microbial concentrations, it lengthens the whole process and compromises the main advantage of the molecular assays, which is the rapid procurement of results. Another strategy is to increase the initial volume of the blood sample (> 1 ml) (21), which has been reported to give promising results and greatly improve the detection rates of PCR-based assays (10) or increase the amount of bacterial DNA, as shown by Trung et al. (22).

We observed 36 samples that were positive according to MP and negative according to BC. This could have been due to the fact that almost half of these samples were obtained from patients on antibiotic treatment, which could explain the negative BC results. The presence of cell-free DNA or non-viable microorganisms or even contaminating DNA could explain the other 19 discordant results (23).

Importantly, MP detected 5 out of the 14 *Candida* spp. and 11 out of the 13 *S. aureus* isolates in our study. Early detection of these two pathogens is essential due to the high mortality rates and risk of haematogenous complications associated with these microbes. In addition, *Candida* requires a longer time for growth in BC than other pathogens, delaying the initiation of treatment by several days. *Candida* spp. was the third most frequently occurring pathogen detected in our study. As expected, most of the positive samples were obtained from ICU patients. We detected a small number of resistance genes. Although the three *mecA* genes detected by MP were confirmed by routine susceptibility testing, 4 methicillin-resistant CoNS were not identified by MP. Furthermore, the only *vanB*-positive *E. faecalis* detected by MP was not isolated by BC. Therefore, the accuracy of MP in detecting resistance genes should be further evaluated.

Despite the many advantages of molecular assays, these tests have several important limitations. First, these methods require specialised equipment and technical experience and are usually expensive. The MP test also has several manual steps that make it laborious and increase the risk of possible contamination. Moreover, its low sensitivity makes its implementation as a routine test difficult in clinical microbiology laboratories. Finally, the sensitivity and specificity of molecular assays vary according to the test, the extraction method, the algorithm used to evaluate the results, the comparative method and the study population. Data published to date support the use of PCR-based tests for specific groups of patients, such as critically ill patients in ICU, those with suspected candidaemia or patients receiving broad-spectrum antibiotics (24,25). There is no consensus about the interpretation of BC-negative and PCR-positive results (low pathogen concentrations, non-viable bacteria and DNAemia, etc.). Further studies are needed to evaluate the role of new molecular assays in the routine microbiological diagnosis of BSI.

In conclusion, sepsis is a time-dependent disease that requires early diagnosis and prompt appropriate treatment to improve prognosis. The MP assay provides results within 6 hours, thereby significantly reducing the amount of time for diagnosis and seems to be especially useful in patients on antimicrobial treatment. Nevertheless, the assay has to be optimised, mainly for greater automation and to facilitate its introduction into routine laboratory workflow. Furthermore, the microbial detection limits of the molecular assay need to be improved, probably by new extraction protocols and greater sample volumes to improve the positive and negative predictive power so that it can be a useful tool in clinical practice, especially regarding its impact on antibiotic use.

## Financial support

This study was supported by the Ministerio de Economía y Competitividad, Instituto de Salud Carlos III, and co-financed by the European Regional Development Fund (ERDF) “A Way to Achieve Europe” and the Spanish Network for Research in Infectious Diseases (REIPI RD12/0015). This study was also supported by grant 2014SGR653 from the Departament d’Universitats, Recerca i Societat de la Informació of the Generalitat de Catalunya, as well as by funding from the Innovative Medicines Initiative (COMBACTE, grant agreement 115523), *Ciber de Enfermedades Respiratorias* (CibeRes CB06/06/0028), 2009 Support to Research Groups of Catalonia 911 and IDIBAPS (CERCA Programme/Generalitat de Catalunya). The funding sources had no role in the design or conduct of the study; collection, management, analysis, or interpretation of the data; preparation, review, or approval of the manuscript; or decision to submit the manuscript for publication.

C.C. is the recipient of a postdoctoral grant (Strategic Plan for Research and Innovation in Health - PERIS 2016-2020). C.M. was working with the IZASA-Werfen Group during the undertaking of this study.

## Acknowledgements

We are indebted to all our medical and nursing colleagues for their assistance and cooperation in this study. All the reagents for the Magicplex™ Sepsis test were provided by the IZASA-Werfen Group.

## Conflicts of interest

The authors declare that they have no conflicts of interest.

## References

1. Ferrer R, Martin-Loeches I, Phillips G, Osborn TM, Townsend S, Dellinger RP, et al. Empiric antibiotic treatment reduces mortality in severe sepsis and septic shock from the first hour: results from a guideline-based performance improvement program. Crit Care Med. 2014 Aug;42(8):1749–55.

2. Kumar A, Roberts D, Wood KE, Light B, Parrillo JE, Sharma S, et al. Duration of hypotension before initiation of effective antimicrobial therapy is the critical determinant of survival in human septic shock. Crit Care Med. 2006 Jun;34(6):1589–96.

3. Candel FJ, Borges Sá M, Belda S, Bou G, Del Pozo JL, Estrada O, et al. Current aspects in sepsis approach. Turning things around. Rev Espanola Quimioter Publicacion Of Soc Espanola Quimioter. 2018 Jun 25;

4. Yagupsky P, Nolte FS. Quantitative aspects of septicemia. Clin Microbiol Rev. 1990 Jul;3(3):269–79.

5. Sinha M, Jupe J, Mack H, Coleman TP, Lawrence SM, Fraley SI. Emerging Technologies for Molecular Diagnosis of Sepsis. Clin Microbiol Rev. 2018 Apr;31(2).

6. Fenollar F, Raoult D. Molecular diagnosis of bloodstream infections caused by non-cultivable bacteria. Int J Antimicrob Agents. 2007 Nov;30 Suppl 1:S7-15.

7. Leitner E, Kessler HH, Spindelboeck W, Hoenigl M, Putz-Bankuti C, Stadlbauer-Köllner V, et al. Comparison of two molecular assays with conventional blood culture for diagnosis of sepsis. J Microbiol Methods. 2013 Mar;92(3):253–5.

8. Grif K, Fille M, Würzner R, Weiss G, Lorenz I, Gruber G, et al. Rapid detection of bloodstream pathogens by real-time PCR in patients with sepsis. Wien Klin Wochenschr. 2012 Apr;124(7–8):266–70.

9. Nieman AE, Savelkoul PHM, Beishuizen A, Henrich B, Lamik B, MacKenzie CR, et al. A prospective multicenter evaluation of direct molecular detection of blood stream infection from a clinical perspective. BMC Infect Dis. 2016 30;16:314.

10. Loonen AJM, Bos MP, van Meerbergen B, Neerken S, Catsburg A, Dobbelaer I, et al. Comparison of pathogen DNA isolation methods from large volumes of whole blood to improve molecular diagnosis of bloodstream infections. PloS One. 2013;8(8):e72349.

11. Wellinghausen N, Kochem A-J, Disqué C, Mühl H, Gebert S, Winter J, et al. Diagnosis of bacteremia in whole-blood samples by use of a commercial universal 16S rRNA gene-based PCR and sequence analysis. J Clin Microbiol. 2009 Sep;47(9):2759–65.

12. Bloos F, Sachse S, Kortgen A, Pletz MW, Lehmann M, Straube E, et al. Evaluation of a polymerase chain reaction assay for pathogen detection in septic patients under routine condition: an observational study. PloS One. 2012;7(9):e46003.

13. Lehmann LE, Hunfeld K-P, Emrich T, Haberhausen G, Wissing H, Hoeft A, et al. A multiplex real-time PCR assay for rapid detection and differentiation of 25 bacterial and fungal pathogens from whole blood samples. Med Microbiol Immunol (Berl). 2008 Sep;197(3):313–24.

14. Fitting C, Parlato M, Adib-Conquy M, Memain N, Philippart F, Misset B, et al. DNAemia detection by multiplex PCR and biomarkers for infection in systemic inflammatory response syndrome patients. PloS One. 2012;7(6):e38916.

15. Kühn C, Disqué C, Mühl H, Orszag P, Stiesch M, Haverich A. Evaluation of commercial universal rRNA gene PCR plus sequencing tests for identification of bacteria and fungi associated with infectious endocarditis. J Clin Microbiol. 2011 Aug;49(8):2919–23.

16. Carrara L, Navarro F, Turbau M, Seres M, Morán I, Quintana I, et al. Molecular diagnosis of bloodstream infections with a new dual-priming oligonucleotide-based multiplex PCR assay. J Med Microbiol. 2013 Nov;62(Pt 11):1673–9.

17. Ljungström L, Enroth H, Claesson BEB, Ovemyr I, Karlsson J, Fröberg B, et al. Clinical evaluation of commercial nucleic acid amplification tests in patients with suspected sepsis. BMC Infect Dis. 2015 Apr 28;15:199.

18. Denina M, Scolfaro C, Colombo S, Calitri C, Garazzino S, Barbui Anna A, et al. Magicplex(TM) Sepsis Real-Time test to improve bloodstream infection diagnostics in children. Eur J Pediatr. 2016;175(8):1107–11.

19. Ziegler I, Fagerström A, Strålin K, Mölling P. Evaluation of a Commercial Multiplex PCR Assay for Detection of Pathogen DNA in Blood from Patients with Suspected Sepsis. PloS One. 2016;11(12):e0167883.

20. Serra J, Rosello E, Figueras C, Pujol M, Peña Y, Céspedes P, et al. Clinical evaluation of the Magicplex Sepsis Real-time Test (Seegene) to detect Candida DNA in pediatric patients. Crit Care. 2012;16(Suppl 3):P42.

21. Hansen WLJ, Bruggeman CA, Wolffs PFG. Evaluation of new preanalysis sample treatment tools and DNA isolation protocols to improve bacterial pathogen detection in whole blood. J Clin Microbiol. 2009 Aug;47(8):2629–31.

22. Trung NT, Hien TTT, Huyen TTT, Quyen DT, Van Son T, Hoan PQ, et al. Enrichment of bacterial DNA for the diagnosis of blood stream infections. BMC Infect Dis. 2016 31;16:235.

23. Opota O, Jaton K, Greub G. Microbial diagnosis of bloodstream infection: towards molecular diagnosis directly from blood. Clin Microbiol Infect Off Publ Eur Soc Clin Microbiol Infect Dis. 2015 Apr;21(4):323–31.

24. Bravo D, Blanquer J, Tormo M, Aguilar G, Borrás R, Solano C, et al. Diagnostic accuracy and potential clinical value of the LightCycler SeptiFast assay in the management of bloodstream infections occurring in neutropenic and critically ill patients. Int J Infect Dis IJID Off Publ Int Soc Infect Dis. 2011 May;15(5):e326-331.

25. von Lilienfeld-Toal M, Lehmann LE, Raadts AD, Hahn-Ast C, Orlopp KS, Marklein G, et al. Utility of a commercially available multiplex real-time PCR assay to detect bacterial and fungal pathogens in febrile neutropenia. J Clin Microbiol. 2009 Aug;47(8):2405–10.

